# Model Adequacy and the Macroevolution of Angiosperm Functional Traits

**DOI:** 10.1101/004002

**Authors:** Matthew W. Pennell, Richard G. FitzJohn, William K. Cornwell, Luke J. Harmon

**Affiliations:** Department of Biological Sciences & Institute for Bioinformatics and Evolutionary Studies, University of Idaho, Moscow, ID 83844 U.S.A.; Department of Biological Sciences, Macquarie University,Sydney, NSW 2109, Australia; School of Biological, Earth and Environmental Sciences, University of New South Wales, Sydney, NSW 2052 Australia

**Keywords:** phylogenetic comparative methods, model adequacy, independent contrasts, Angiosperm functional traits

## Abstract

Making meaningful inferences from phylogenetic comparative data requires a meaningful model of trait evolution. It is thus important to determine whether the model is appropriate for the data and the question being addressed. One way to assess this is to ask whether the model provides a good statistical explanation for the variation in the data. To date, researchers have focused primarily on the explanatory power of a model relative to alternative models. Methods have been developed to assess the adequacy, or absolute explanatory power, of phylogenetic trait models but these have been restricted to specific models or questions. Here we present a general statistical framework for assessing the adequacy of phylogenetic trait models. We use our approach to evaluate the statistical performance of commonly used trait models on 337 comparative datasets covering three key Angiosperm functional traits. In general, the models we tested often provided poor statistical explanations for the evolution of these traits. This was true for many different groups and at many different scales. Whether such statistical inadequacy will qualitatively alter inferences draw from comparative datasets will depend on the context. Regardless, assessing model adequacy can provide interesting biological insights — how and why a model fails to describe variation in a dataset gives us clues about what evolutionary processes may have driven trait evolution across time.

## Introduction

A statistical model may provide the best explanation for a dataset compared to a few other models but still be a very poor explanation in terms of capturing the patterns of variation present in the data. For simple linear regression models, absolute model fit, or adequacy, is commonly assessed by simply plotting the data 45 alongside the best regression line. While not quantitative, visualizing the bivariate distribution can provide important insights regarding the fit of the model that are not captured by summaries such as the *R^2^* or *p*–value, such as whether the relationship is indeed linear (for a classic case study, see Anscombe, 1973). For these types of models, there are also a wide variety of statistical tests of model adequacy (e.g., 50 the relationship between the residuals and the independent variable, χ^2^ goodness-of-fit test, etc.) that compliment our visual intuition about model adequacy. Such formal tests used alongside informal visualizations can help researchers assess whether the inferences drawn from the fitted model are meaningful and, more interestingly, suggest how a model can be improved (Gelman and Shalizi, 2013).

Modern phylogenetic comparative methods for investigating trait evolution are almost exclusively model-based (recently reviewed in O’Meara, 2012; Pennell and Harmon, 2013), meaning that inferences are contingent on both the phylogenetic tree and the model for the traits. Selecting a good model is therefore essential for making robust inferences. Researchers typically use likelihood ratio tests or 60 Information Theoretic measures (i.e., AIC, BIC) to select amongst models (Mooers et al., 1999; Harmon et al., 2010; Hunt, 2012) but these only provide a measure of relative fit. Unlike in linear regression models, for most phylogenetic models of trait evolution, it usually very challenging to visually assess the adequacy of a model. This problem is compounded for relatively complex models such as multi-65 rate Brownian motion (O’Meara et al., 2006; Eastman et al., 2011) or multi-optima Ornstein-Uhlenbeck models (Hansen, 1997; Butler and King, 2004; Beaulieu et al., 2012; Uyeda and Harmon, 2014). One can plot the trait values at the tips of the phylogeny but determining “by eye” whether this distribution is consistent with the traits having evolved under the proposed model is difficult at small scales and 70 impossible for large phylogenies.

A number of statistical procedures have been proposed to quantitatively assess the absolute fit of a model of trait evolution (e.g., Garland et al., 1992, Garland et al., 1993; Purvis and Rambaut, 1995; Díaz-Uriarte and Garland, 1996; Freckleton and Harvey, 2006; Boettiger et al., 2012; Slater and Pennell, 2014; Beaulieu et al., 2013; Blackmon and Demuth, 2014). These can be generally classified into two types of approaches. The first are tests for specific deviations from a particular model. In the early days of phylogenetic comparative biology, the focus was primarily on inferring character correlations in order to test hypotheses regarding adaptation (e.g., Felsenstein, 1985; Grafen, 1989; Harvey and Pagel, 1991; Lynch, 1991). Accordingly, a number 80 of tests were developed to assess the reliability of assuming a Brownian motion (BM) model, which formed the basis for all phylogenetic tests of continuous character evolution at the time. Garland et al. (1992) proposed plotting the standardized independent contrasts (*sensu* Felsenstein, 1985) against the standard deviation of each contrast. If the contrasts and their standard deviations are correlated, this would suggest that the model (or the phylogeny) is not adequate. Purvis and Rambaut (1995) suggested using the relationship between the contrasts and the height above the root at which they were generated (see also Freckleton and Harvey, 2006, for a slight modification of this test). Similarly, Beaulieu et al. (2013) and Blackmon and Demuth (2014) used summary statistics to evaluate whether a set of discrete character data was consistent with some variant of a Mk model (Pagel, 1994). These are all very useful ideas, and we have adopted many of these in the method we present below, but each approach is only informative with respect to a single type of misspecification for a single type of model.

The second class of approaches is to use Monte Carlo simulations to compare an observed dataset to those expected under a model. Garland et al. (1993) and Díaz-Uriarte and Garland (1996) developed such an approach two decades ago. However, as this work preceded the development of analytical tools for fitting alternative (i.e., non-BM) models, the simulation parameters were not estimated directly from the data and therefore “reasonable” parameter estimates had to be chosen *a priori*. More recently, two approaches have been suggested for assessing model adequacy using parameters estimated directly from the data. Boettiger et al. (2012) proposed simulating data under two candidate models using the maximum likelihood parameter estimates from each model and then fitting both models to each simulated dataset. Under each of the two simulation conditions, they calculated the likelihood ratio; after many simulations, a distribution of likelihood ratios could be obtained for each ease and these distributions compared to assess whether there was sufficient information in the data to favor one model over the other. Slater and Pennell (2014) used posterior predictive simulation (explained below) to assess the absolute fit of an “early burst” model of trait evolution, in which rates of trait evolution declined through time, compared to that of a BM model. Both Boettiger et al. (2012) and Slater and Pennell (2014) focused on the ability to distinguish between two models using absolute fit. Our aim here is more general: we want to compare the fit of the model to the universe of possible models.

In this paper, we propose a statistical framework for assessing the adequacy of phylogenetic models of quantitative trait evolution that generalizes previous approaches to a wide variety of alternative models. Our central thesis is that assessing model adequacy in a general way can provide valuable insights into evolutionary processes and patterns that are not evident from comparing a limited set of models. For example, one common application of phylogenetic trait models is to make inferences regarding the rate (tempo) of evolution using model selection (e.g., Mooers et al., 1999; Harmon et al., 2010; Hunt, 2012; Slater, 2013). Statements about rates are only informative in the context of a specific model (Hunt, 2012). It is therefore imperative to know if a model is really capturing the variation of the data in absolute terms. In an oft-cited example of this model comparison approach, Harmon et al. (2010) compared three simple models of trait evolution across 49 clades and tallied the frequency with which the models were prefered in order to draw inferences about general patterns. We perform the same analysis but on a much larger scale. We analyze 337 datasets on three important Angiosperm (flowering plants) functional traits using a recently published time-calibrated phylogeny (Zanne et al., 2014). We then assess the adequacy of the best-fitting model across all the datasets to determine how often one of these simple models would be adequate to make reliable inferences about rate of trait evolution.

## A general framework for assessing the adequacy of phylogenetic models

We focus here on models that describe the evolution of a single, continuously valued trait. More specifically, our approach works for models that predict that trait values at the tips come from a multivariate normal distribution. This applies to most models of quantitative trait evolution that have been developed to date (see below for details on the scope of the method).

If we have a phylogenetic tree consisting of *n* lineages and data on the trait values observed at each tip *X* (*X* = *x*_1_, *x*_2_,…, *x_n_*), we can fit a model 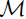 with parameters *θ* to describe the pattern of trait evolution along the phylogeny. There are two primary ways of fitting models to comparative data. The first is use to obtain a point estimate of 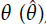 via maximum likelihood (ML), restricted maximum likelihood (REML), least-squares, etc. The second is to estimate the posterior probability distribution 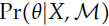 using Bayesian approaches. For the models used in comparative biology, estimating 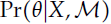 requires using Markov chain Monte Carlo (MCMC) machinery to sample values of *θ*.

Most analyses using comparative data aim to answer one of the following questions: what values of *θ* best explain *X* given 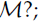 or, does 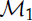 explain the data better than 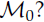 Our approach is conceptually distinct in that we want to ask, how likely is it that model 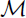 with parameters *θ* would produce a dataset similar to *X* if we re-ran evolution?

While optimizing and Bayesian approaches to model-fitting are philosophically different from one another, our approach to assessing model adequacy is the same for both: (1) fit the model of trait evolution; (2) rescale the branch lengths of the phylogeny to place the data on a standard scale; (3) calculate a set of test statitics, 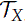 which provide statistical summmaries of the observed data; (4) simulate many new datasets 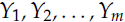 under the model using the estimated parameters; (5) calculate test statistics on the simulated data 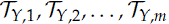(6) compare 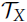 to the distribution of 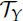. If 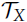 deviates significantly from the distribution of 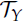, we can consider the model as an inadequate descriptor (see figure 1).

**Figure 1.**
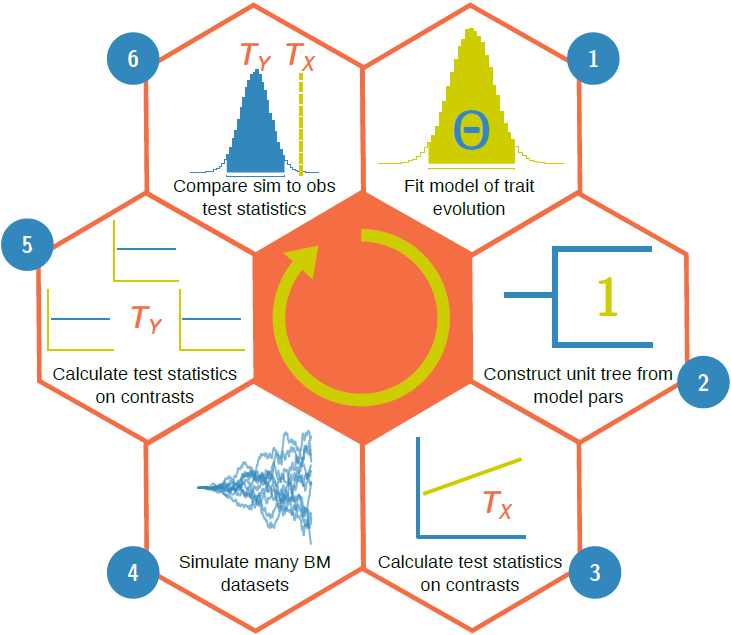
Schematic diagram representing our approach for assessing model adequacy. (1) Fit a model of trait evolution to the data; (2) use the estimated model parameters to build a unit tree; (3) compute the contrasts from the data on the unit tree and calculate a set of test statistics 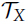; (4) simulate a large number of datasets on the unit tree, using a BM model with *σ*^2^ = 1; (5) calculate the test statistics on the contrasts of each simulated dataset 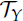 and (6) compare the observed and simulated test statistics. If the observed test statistic lies in the tails of the distribution of simulated test statistics the model can be rejected as inadequate. The rotational circle in the center of the diagram indicates that assessing model adequacy is an iterative process. If a model is rejected as inadequate, the next step is to propose a new model and repeat the procedure.

If we have a point estimate of the model parameters, we simulate 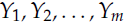 on the phylogeny according to 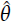 and 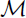. We then compare a single set of test statistics 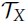 calculated from our observed data to the distribution of values for 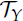 computed across all *m* simulated datasets. In statistical terminology, this procedure is known as parametric bootstrapping. Parametric bootstrapping is likely familiar to phylogenetic biologists in the form of the Goldman–Cox test (Goldman, 1993) for assessing the adequacy of sequence evolution models and more recently, the phylogenetic Monte Carlo approach of Boettiger et al. (2012).

If we have a posterior probability distribution 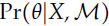, we can assess model adequacy using posterior predictive simulation (Rubin, 1984; Gelman et al., 1996). We obtain new datasets by sampling from a second distribution, the posterior predictive distribution

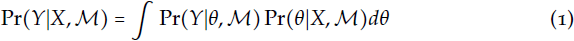
where 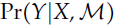 is the probability of a new dataset 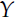 given *X* and 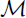 averaged over the posterior distribution of the parameters. (Sampling from 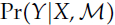 is equivalent to simulating datasets using paramaters drawn from the posterior distribution.) Therefore, the datasets 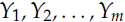 previously developed for models in molecular phylogenetics (Bollback, 2002; Reid et al., 2014; Lewis et al., 2014; Brown, 2014), and recently for PCMs (Slater and Pennell, 2014), but have not been widely adopted in either field.

## Test statistics

No simulated dataset will ever be exactly the same as our observed dataset. We therefore need to choose informative test statistics in order to evaluate whether the model predicts datasets that are similar to our observed dataset in meaningful ways. As the states at the tips of the phylogeny are not independent — this is why we are using PCMs in the first place! — calculating test statistics on the data directly is not generally informative for models in comparative biology. We account for the non-independence of the observed data by calculating test statistics on the set of contrasts (i.e., “phylogenetically independent contrasts“; Felsenstein, 1985) computed at each node. (We refer readers to Felsenstein, 1985; Rohlf, 2001; Blomberg et al., 2012, for details on how contrasts are calculated.) Under Brownian motion (BM) the contrasts will be independent and identically distributed (i.i.d.) according to a normal distribution with mean 0 and standard deviation *σ* (i.e., contrasts are 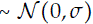), where *σ*^2^ is the BM rate parameter (Felsenstein, 1985). This i.i.d. condition allows us to perform standard statistical tests on the contrasts.

The choice of what test statistics to use for assessing model adequacy is ultimately one of balancing statistical intuition and computational effort. We have chosen the following set of six test statistics to compute on the contrasts because they capture a range of possible model violations and have well-understood statistical properties. All of these essentially evaluate whether the contrasts come from the distribution expected under BM.

*M*_SIG_ The mean of the squared contrasts. This is equivalent to the REML estimator of the Brownian motion rate parameter *σ*_2_ (Garland et al., 1992; Rohlf, 2001). *M*_SIG_ is a metric of overall rate. Violations detected by *M*_SIG_ indicate whether the overall rate of trait evolution is over-or underestimated.

*C*_VAR_ The coefficient of variation (standard deviation/mean) of the absolute value of the contrasts. If *C*_VAR_ calculated from the observed contrasts is greater than that calculated from the simulated contrasts, it suggests that we are not properly accounting for rate heterogeneity across the phylogeny. If CVAR from the observed is smaller, it suggests that contrasts are more even than the model assumes. We use the coefficient of variation rather than the variance because the mean and variance of contrasts can be highly correlated.

*S*_VAR_ The slope of a linear model fit to the absolute value of the contrasts against their expected variances (following Garland et al., 1992). Each (standardized) contrast has an expected variance proportional to the sum of the branch lengths connecting the node at which it is computed to its daughter lineages (Felsenstein, 1985). Under a model of BM, we expect no relationship between the contrasts and their variances. We use *S*_VAR_ to test if contrasts are larger or smaller than we expect based on their branch lengths. If, for example, more evolution occurred per unit time on short branches than long branches, we would observe a negative slope. If *S*_VAR_ calculated from the observed data deviates substantially from the expectations, a likely explanation is branch length error in the phylogenetic tree.

*S*_ASR_ The slope of a linear model fit to the absolute value of the contrasts against the ancestral state inferred at the corresponding node. We estimated the ancestral state using the least—squares method suggested by Felsenstein (1985) for the calculation of contrasts. (We note that this is not technically an ancestral state reconstruction [see Felsenstein, 1985]; it is more properly thought of as a weighted average value for each node.) We used this statistic to evaluate whether there is variation in rates relative to the trait value. For example, do larger organisms evolve proportionally faster than smaller ones?

*S*_HGT_ The slope of a linear model fit to the absolute value of the contrasts against node depth (after Purvis and Rambaut, 1995). This is used to capture variation relative to time. It is alternatively known as the “node-height test” and has been used to detect early bursts of trait evolution during adaptive radiations (see Freckleton and Harvey, 2006; Slater and Pennell, 2014, for uses and modifications of this test).

*D*_CDF_ The D–statistic obtained from Kolmolgorov-Smirnov test from comparing the distribution of contrasts to that of a normal distribution with mean 0 and standard deviation equal to the root of the mean of squared contrasts (the expected distribution of the contrasts under BM; see Felsenstein, 1985; Rohlf, 2001). We chose this to capture deviations from normality. For example, if traits evolved via a “jump–diffusion” type process (Landis et al., 2013), in which there were occasional bursts of rapid phenotypic evolution (Pennell et al., 2013), the tip data would no longer be multivariate normal owing to a few contrasts throughout the tree being much larger than the rest (i.e., the distribution of contrasts would have heavy tails).

Alternative test statisics are certainly possible. One could, for instance, calculate the median of the squared contrasts, the skew of the distribution of contrasts, etc. If the generating model was known, we could use established procedures for selecting a set of sufficient (or, approximately sufficient; Joyce and Majoram, 2008) test statistics for that model, as is typically done when computing likelihood ratio tests. However, the aim of our approach is assess the fit of a proposed model without reference to a true model. Our test statistics will detect many types of model misspecification but this does not mean that they will necessarily detect every type of model misspecification. We encourage researchers interested in specific questions to explore alternative test statistics that capture deviations relevant to the problem at hand.

An additional challenge is determining how to deal with the statistical problems (i.e., inflated Type-1 error rates) that may be introduced when using many test statistics. In our analyses, we chose not to correct our p–values for multiple comparisons (using Bonferroni, false discovery rates, etc.). We did this for a number of reasons. First, our tests are not truly independent and the degree of correlation between test statistics will necessarily depend on the “true” model of trait evolution. Second, as argued by Gelman (2006), we might be interested in the specific aspects of the data that different from the expectations under the model; rather than focus on whether a model should be accepted or rejected, we “want to understand the limits of its applicability in realistic replications” (p. 175).

## Beyond Brownian motion

All of our test statistics are designed to evaluate the adequacy of a BM model of trait evolution. However, if we propose a different model for the evolution of the trait, such as an Ornstein—Uhlenbeck (OU; Hansen, 1997) process, then the expected distribution of the contrasts is different. For example, under an OU model, contrasts will not be i.i.d. (Hansen, 1997). The expected distribution of contrasts under most models of trait evolution, aside from BM, is not formally characterized and even if it was, this would necessitate a specific set of test statistics for every model proposed.

Our solution to this problem is to create what we term a “unit tree”, which is a phylogenetic tree transformation that captures the dynamics of trait change under a particular evolutionary model. For a particular evolutionary model 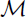(with parameter values *θ*), we define a unit tree as a phylogenetic tree that has the following property: the length of branch *b*, *ν_b_*, is equal to the amount of variance expected to accumulate over *i* under 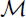, *θ*. The variance is standardized, such that the expected distribution of the trait data on the unit tree is equal to that of a Brownian Motion (BM) model with a rate *σ*^2^ equal to 1.

If the fitted model is adequate, the trait data at the tips of the unit tree will have the same distribution as data generated under a BM process with a rate of 1 and the contrasts will be distributed according to a standard normal distribution (hence the name, unit tree). Creating the unit tree from the estimated model parameters prior to computing the contrasts generalizes the test statistics to most models of quantitative trait evolution (but see Landis et al., 2013; Schraiber and Landis, 2014, for exceptions).

We also emphasize that because the contrasts are calculated on the unit tree, the test statistics all must depend on both the data and the model; for this reason, the Bayesian version of our approach produces a distribution of observed test statistics. Once we have created the unit tree from the estimated parameters, new datasets can be simulated under the model simply using a BM process with *σ*^2^ = 1, which has the added benefit of being computationally efficient. The distribution of test statistics calculated on these simulated data sets can then be compared to the test statistics from the observed data.

## Details of unit tree construction and the scope of this approach

Here we formalize our definition of the unit tree and delimit the scope of our approach. Readers can skip this section without missing the main point. A unit tree can be constructed from any evolutionary model where the trait has expected variance-covariance matrix **V** that satisfies the (generalized) 3-point condition proposed by Ho and Ané (2014) and the data follows a multivariate normal distribution. A matrix **V** has a strict 3-point structure if the following condition holds: for any lineages *i, j, k*, the two smallest of *V_ij_*, *V_ik_*, *V_jk_* are equal. Under a simple BM model it is straightforward to show that this condition holds. If **C** is the matrix representation of the phylogeny (such that *C_ij_* is the shared path length between lineages *i* and *j*), then by the nature of the tree structure, the 3-point condition will hold for **C**. Since under BM **V** = *σ^2^***C**, then **V** will also be 3-point structured. The same holds true for any evolutionary model that is a branch length transformation of a BM model including the λ, δ, *κ* models (Pagel, 1997, Pagel, 1999) and models where rates change through time (the “Early Burst” or EB model, also referred to as the Accelerating/Decelerating Change, ACDC, model; Blomberg et al., 2003; Harmon et al., 2010) or across the tree (O’Meara et al., 2006; Thomas et al., 2006; Eastman et al., 2011; Revell et al., 2012; Thomas and Freckleton, 2012). Standard error can be incorporated into any of these models by simply adding a species-specific scalar to each element of the diagonal. For all of the models where the 3-point condition applies, we can construct a unit tree by setting

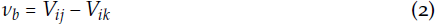

where *V_ij_* and *V_jk_* are, by the requirements of the 3-point structured condition, equal to one another. Once all branches have been transformed, the contrasts computed on the unit tree will be i.i.d. 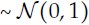 under the model in question.

The OU model of trait evolution also generates 3-point structured matrices when the tree is ultrametric; this is true of both single optimum and multi-optima models (Ho and Ané, 2014). However, while the variance structure can easily be transformed to a BM-like tree, the contrasts on this tree will not necessarily be distributed according to a standard normal. For example, while it is often assumed when fitting a single regime OU model that the ancestor is at the optimum trait value (see, for example Harmon et al., 2010), this need not be the case. Furthermore, if there are multiple optima on the phylogeny (Hansen, 1997; Butler and King, 2004; Ingram and Mahler, 2013; Uyeda and Harmon, 2014), lineages will necessarily be tracking optima that are different from the root state. Therefore, a transformation must also be made to the data in addition to the branch lengths of the phylogeny to produce contrasts that have are i.i.d. according to a standard normal.

To accomplish this, we again turn to the recent work of Ho and Ané (2014). In addition to 3-point structured matrices, Ho and Ané defined a broader condition: a matrix of the form

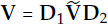

is considered to have a generalized 3–point structure if 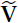 is 3-point structured and **D**_1_ and **D**_2_ are diagonal matrices. Ho and Ané (2014) prove that many phylogenetic models are indeed of this class, including multi-optimum OU models (Butler and King, 2004; Ingram and Mahler, 2013; Uyeda and Harmon, 2014), those with varying rates and models across the tree (e.g., Beaulieu et al., 2012) as well as to OU models fit to non-ultrametric trees. For any model that satisfies the generalized 3-point condition and where the data is assumed to come from a multivariate normal distribution, there exists some transformation to the tree (appling Equation 2 to 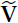) and data (using **D**_1_ and **D**_2_) that will produce a unit tree with standard normal contrasts. We note that Slater (2014) recently pointed out that for OU models fit to non-ultrametric trees, there is no valid transformation that can make **V** BM-like. While this is indeed correct, it is however, possible to get a BM-like tree by adding a species-specific scalar to the data matrix (Ho and Ané, 2014). Therefore, once the proper tree and data transformations have been made, all the test statistics described above can apply.

The above also applies to phylogenetic regression models (Grafen, 1989; Lynch, 1991; Martins and Hansen, 1997) of the form

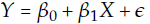

In these models, the error variance is structured by phylogeny assuming some model of trait evolution such that 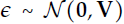. In these regression models **V** represents the variance-covariance matrix of the residuals rather than the traits (Rohlf, 2001). Therefore if **V** is either 3—point or generalized 3—point structured, the tree (and possibly data) can be transformed such that the contrasts on the residuals will be i.i.d. standard normal. This fact allows researchers to use our approach to assess the adequacy of a trait model for understanding correlations between traits. We note however that as **V** only affects the error structure for these models, alternative approaches (see for example Gelman et al., 2003, ch. 6) will be required to assess the adequacy of the mean structure 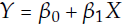 of the model.

## Simulations

As a verification of our method, we conducted a brief simulation study. We focused here on assessing Type—1 error rates. As above, we emphasize that these are not necessarily the most important quantities when thinking about model adequacy, but they do provide a useful metric for demonstrating that our code is functioning correctly. The philosophy behind approaches such as ours is that the “true” model is outside of the candidate set. We want to ask whether a given model can adequately describe the variaton in the data. If it does, we can consider it statistically adequate even if it is not the true model or even the best model in our set (see Discussion for comments on the relationship between model adequacy and model selection). Furthermore, while it is certainly interesting to examine what types of deviations in model space produce what types of deviations in the various test statistics, the number of possible simulation conditions is infinitely broad.

We simulated data under BM, single–optimum OU, and EB (the same models we used in the analysis; see below). For each set of simulating conditions, we simulated trees of 50, 100, and 200 taxa under a pure-birth process, then rescaled the tree to be unit height. For BM, we set *σ*^2^ = 1. For OU, we used *σ*^2^ = 1 but varied the “selection” parameter *α(α =*{1,2,4}). For EB, we again set *σ*^2^ to be 1 and varied *a*, the exponential rate of decline (see Harmon et al., 2010; Slater and Pennell, 2014, for details), was set to be *a =*{log(0.01),log(0.02),log(0.04)}. For each parameter combination, we ran 500 simulations under two sets of conditions: (1) assuming no measurement error; and (2) assuming known error rates of 5% of the expected variance in trait values across the phylogeny. We then fit the corresponding model using maximum likelihood and evaluated the Type-1 error under each set of conditions. All simulations were conducted using diversitree (FitzJohn, 2012).

## The adequacy of models for the evolution of plant functional traits

### Data

We used a phylogeny of Angiosperms, containing 30,535 species, from a recent study by Zanne et al. (2014). We conducted all analyses on the MLE of the phylogeny (available on DRYAD, doi:10.5061/dryad.63q27/3). We used existing large datasets on three functionally important plant traits: specific leaf area (SLA, defined as fresh area/dry mass), seed mass, and leaf nitrogen content (% mass). Seed mass is a crucial part of species' life-history strategy (Leishman et al., 2000; Westoby et al., 2002) and SLA and leaf nitrogen content are important and widely measured components of species' carbon capture strategies (Wright et al., 2004). Understanding the macroevolutionary patterns of these three traits can provide key insights into the evolutionary processes that have shaped much of plant diversity (Cornwell et al., 2014). The SLA and leaf nitrogen data comes from Wright et al. (2004) with additional SLA data from the LEDA project (Kleyer et al., 2008). Seed mass data comes the Kew database (Royal Botanical Gardens, Kew, 2014). We used an approximate grepping approach to find and correct spelling mistakes and synonymy tools from The Plant List (2014) to match the trait databases to the Zanne et al. phylogeny. The full data set includes 3293 species for SLA, of which 2200 match species in the Zanne et al. (2014) tree. For seed mass, the dataset included 22,817 species with 11,107 matched the phylogeny. For leaf nitrogen content, we have data for 1574 species with 936 included in the tree. See https://github/richfitz/modeladequacy for specific locations and scripts to access and process the original data.

We log—transformed all data prior to analysis. We did this for biological reasons rather than to conform the data to the assumptions of the model (Houle et al., 2011). It is more meaningful to model trait evolution as a multiplicative process rather than an arithmetic one. An increase of two grams is much more significant for the seed of an orchid than the seed of a palm tree. However, we should recognize that both of these rationales are essentially statements about model adequacy and thus the validity of the log transformation can be quantitatively assessed. We ran our analysis on both the raw and log transformed data.

Because the vast majority of the species are only represented by a single record, it was not possible to use a species—specific estimate of trait standard error (SE) to account for either measurement error or intraspecific variation. As an alternative, we estimated a single SE for each trait by calculating the mean standard deviation for all species for which we had multiple measurements. The assumption of a constant SE across all species is unlikely to be correct, but even an inaccurate estimate of error is better than assuming none at all (Hansen and Bartoszek, 2012).

### Analysis

We first matched our trait data to the whole phylogeny and then extracted sub-clades from this dataset in a three ways: (1) by family; (2) by order; and (3) by cutting the tree at 50 My intervals and extracting the most inclusive clades (named or unnamed) for which the most recent common ancestor of a group was younger than the time-slice. (The crown age of Angiosperms is estimated to be ~243 my in the MLE tree and the tree was cut at 50, 100, 150, and 200my.) We kept only sub-clades for which there was at least 20 species present in both the phylogeny and trait data so that we had a reasonable ability to estimate parameters and distinguish between models (Boettiger et al., 2012; Slater and Pennell, 2014). For SLA, this left us with 72 clades, seed mass, 226 clades, and leaf nitrogen content, 39 clades (337 in total). We note that these datasets are not independent as many of the same taxa were included in family, order and multiple time—slice subtrees.

Following Harmon et al. (2010), we considered three simple models of trait evolution: (1) BM, which can be associated with genetic drift (Lande, 1976; Felsenstein, 1988; Lynch and Hill, 1986; Lynch, 1990; Hansen and Martins, 1996), randomly–varying selection (Felsenstein, 1973), or the summation of many independent processes over macroevolutionary time (Hansen and Martins, 1996; Uyeda et al., 2011; Pennell et al., 2013); (2) single optimum OU, which is often assumed to represent stabilizing selection (following Lande, 1976), though we think a more meaningful interpretation is that it represents an “adaptive zone” (Hansen, 2012; Pennell and Harmon, 2013); and (3) EB, which was developed as a phenomenological representation of a niche-filling process during an adaptive radiation (Blomberg et al., 2003; Harmon et al., 2010). We fit each of these models to all 337 subclades in our dataset. We then used the approach we developed to assess the adequacy of each fitted model.

All of the analyses conducted in this paper were conducted using both likelihood and Bayesian inference. We did so to demonstrate the scope of our approach and because both ML and Bayesian inference are commonly used in comparative biology. We emphasize that our approach is not tied to any single statistical paradigm.

For the likelihood analyses, we fit the three models (BM, OU, and EB) using ML with the diversitree package (FitzJohn, 2012). We calculated the AIC score for each model. We then constructed a unit tree for each subtree, trait and model combination using the maximum likelihood estimates of the parameters. We calculated the six test statistics described above (*M*_SIG_, *C*_VAR_, *S*_VAR_, *S*_ASR_, *S*_HGT_, *D*_CDF_) on the contrasts of the data. We simulated 1000 datasets on each unit tree using a BM model with *σ*^2^ = 1 and calculated the test statistics on the contrasts of each simulated data set.

For the Bayesian analysis, we fit the same models as above using a MCMC approach, sampling parameter values using slice sampling (Neal, 2003), as implemented in `diversitree` (FitzJohn, 2012). For the BM model we set a broad uniform prior on 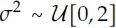, the upper bound being substantially larger than the ML estimate of o^2^ for any clade. For the OU model, we used the same prior for *σ*^2^ and drew *α* values, the strength of attraction to the optimum, from a Lognormal(*μ* = log(0.5), *σ* = log(1.5)) distribution. A complication involved in fitting OU models is deciding what assumptions to make about the state at the root *Z*_0_. Here, we follow other authors (Butler and King, 2004; Harmon et al., 2010) and assume that *Z*_0_ is at the optimum. For the EB model, we again used the same prior for *σ*^2^ and a uniform prior on a, the exponential rate of decrease in *σ*^2^, such that 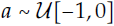 (the minimum value is much more negative than we would typically expect; Slater and Pennell, 2014).

Again, for each model/trait/subtree combination, we ran a Markov chain for 10,000 generations. Preliminary investigations demonstrated that this was more than sufficient to obtain convergence and proper mixing for these simple models. After removing a burn-in of 1000 generations, we calculated the Deviance Information Criterion (DIC, a Bayesian analog of AIC; Spiegelhalter et al., 2002) for each model. We drew 1000 samples from the joint posterior distribution. For each of the sampled parameter sets, we used the parameter values to construct a unit tree and calculated our six test statistics on the contrasts. We then simulated a dataset on the same unit tree and calculated the test statistics on the contrasts of the simulated data.

In the likelihood analyses, for each dataset, we had one set 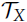 of observed test statistics and a 1000 sets 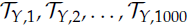 of test statistics calculated on data simulated on the same unit tree. In the Bayesian version, we had 1000 sets of observed test statistics 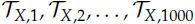 using a different unit tree for each set and 1000 sets of simulated test statistics 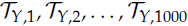, each 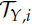 corresponding to the unit tree used to compute 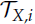.

For both types of analyses, we report two–tailed *p*–values (i.e., the probability that the observed that a simulated test statistic was more extreme than the observed). As a multivariate measure of model adequacy, we calculated the Mahalanobis distance, a scale-invariant metric, between the observed test statistics and the mean of our simulated test statistics, taking into account the covariance structure between the simulated test statistics. We took the log of the KS D-statistic, DCDF, as the Mahanalobis measure assumes data is multivariate normal and the D-statistic is bounded between 0 and 1. For the Bayesian analyses, we report the mean of the distribution of Mahalanobis distances. All analyses were conducted in R v3.0.2 (R Development Core Team, 2013). Scripts to fully reproduce all analyses are available at https://github.com/richfitz/modeladequacy.

### A case study: seed mass evolution in the Meliaceae and Fagaceae

As an illustration of our approach, we present a case study examining seed mass evolution in two tree families, the Meliaceae, the “mahogany family”, and Fagaceae, which contains oaks, chestnuts and beech trees. The trait data and phylogeny for both groups are subsets of the larger dataset used in the analysis. Superficially, these datasets are quite similar. Both are of similar size (Meliaceae: 44 species in the dataset, 550 in the clade; Fagaceae: 70 species in the dataset and 600 in the clade), age (crown age of Meliaceae: ~53my; Fagaceae: ~40my) and are ecologically comparable in terms of dispersal strategy and climatic niche.

As described above, we fit three simple models of trait evolution (BM, OU, EB) to both datasets using ML and computed AIC weights (AIC_*w*_; Akaike, 1974; Burnham and Anderson, 2004) for the three models. For both datasets, an OU model was overwhelmingly supported (AIC_*w*_ > 0.97 for both groups). Therefore, looking only at relative model support, we might conclude that similar evolutionary processes are important in these two clades of trees.

Examining model adequacy provides a different perspective. We took the MLE of the parameters from the OU models for each dataset and constructed a unit tree based on those parameters. We calculated our six test statistics on the contrasts of the data, then simulated 1000 datasets on the unit tree and calculated the test statistics on the contrasts of each simulated dataset ((Figure 2). For seed mass evolution in Meliaceae, the OU model was an adequate model; all six observed test statistics were in the middle of the distribution of simulated test statistics (*M*_SIG_: *p* = 0.921, *C*_VAR_: *p* = 0.605, *S*_VAR_: *p* = 0.979, *S*_ASR_: *p* = 0.485, *S*_HGT_: *p* = 0.170, *D*_CDF_: *p* = 0.657). In contrast, for Fagaceae we found that the test statistics calculated with an OU model lay outside the expected values for *S*_VAR_ (p ≈ 0) and *S*_HGT_ (p = 0.014) suggesting that the process of evolution that gave rise to this data was more complex that that captured by a simple OU process. Specifically, we would infer that rates of evolution depend on the length of the branches (*S*_VAR_), which may indicate phylogenetic error, and that the model is failing to fully capture variation through time (*S*_HGT_. The rest of the observed test statistics did not differ significantly from the simulated test statistics (*M*_SIG_: *p* = 0.298, *C*_VAR_: *p* = 0.837, *S*_ASR_: *p* = 0.074, *D*_CDF_: *p* = 0.551). This example illustrates the distinction between the conventional approach to model selection in PCMs and model adequacy. Selecting amongst a limited pool of models does not give a complete picture of the amount of variation that a chosen model is actually capturing.

**Figure 2.**
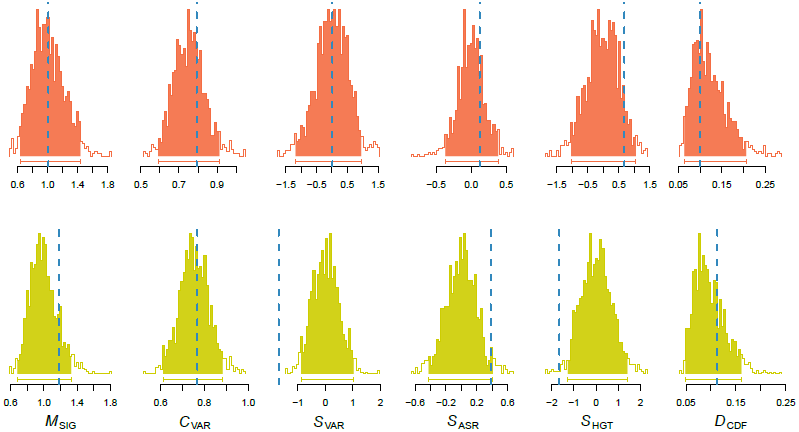
Illustration of our approach to model adequacy. We fit three models (BM, OU, and EB) to seed mass data from two different tree families, the Meliaceae (top panel, red) and the Fagaceae (bottom panel, yellow). In both cases, an OU model (analyzed here) was strongly supported when fit with ML. The plotted distributions are the test statistics (*M*_SIG_, *C*_VAR_, *S*_VAR_, *S*_ASR_, *S*_HGT_, *D*_CDF_) calculated from the contrasts of the simulated data; the bars underneath the plots represent 95% of the density. The dashed vertical lines are the values of the test statistics calculated on the contrasts of the observed data. Using our test statistics, an OU model appears to be an adequate model for the evolution of seed mass in the Meliaceae; for all of the test statistics, the observed test statistic lies in the middle of the distribution of simulated test statistics. For the Fagaceae, the slopes of the contrasts against their expected variances *S*_VAR_ and node height *S*_HGT_ are much lower than the expectations under the model.

## Results

### Simulations

In our simulations, we found that when we assessed the adequacy of the generating model, all of the test statistics showed Type-1 errors that were consistently around or less than 0.05. This was true across models, parameters, tree sizes and did not depend on whether we included a known SE or not (Figure S7, Figure S8, and Figure S9). These results demonstrate that our unit tree construction is working properly; if the MLE is equal to the generating value, then the constrasts will be i.i.d. 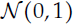 and standard normal statistical properties will apply. Some of the test statistics are very conservative (have very low Type-1 error rates) under some models. We are not aware of any general statistical theory that will allow us to predict the conditions under which a test statistic will have low power to detect deviations from the expected distributions. However, there is an intiutive explanation for this pattern. Consider for example, our test statistic *M*_SIG_. As mentioned above, this is equivalent to the REML estimate of *σ*^2^. When we fit BM (or, a more general model, of which BM is a special case), and then rescale the tree with 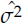, the observed contrasts on the unit tree will effectively be minimized with respect to this quantity and all of the contrasts on the simulated dataset will tend to be larger than our observed contrasts. So if the quantity captured by the test statistic is tightly correlated with one of the parameters being optimized in the model, this test statistic will tend to have low power to detect deviations from this model.

We also found that by using multiple test statistics and reporting a Type-1 error if any of the test statistics deviated significantly from expectations, the error rate increased substantially (up to around 20% under some conditions). However, as we discuss above, we do not think that this is necessarily a defect of the analysis and are not overly concerned with this error rate. Looking at what test statistics were violated and how they were violated is much more meaningful than simply rejecting or accepting a model based on the overall *p*–value. Furthermore, the degree to which the Type-1 error rate will rise with multiple comparisons will be a complex function of the generating model and the size of the dataset and there is no suitable general correction that we know of.

### Models for the evolution of Angiosperm functional traits

Our results for likelihood and Bayesian inference were broadly similar; for conciseness, we present only the results from the likelihood analyses here. Results from the Bayesian analysis are presented in the Supplemental Material. Full results from all analyses can be reproduced using code and workflows available at https://github.com/richfitz/modeladequacy.

Across the 337 subclades, we found widespread support for OU models. For 236 of clades, OU had the highest AIC_*w*_. OU had ~100% of the AIC_*w*_ in 27 clades and >75% of the weight in 189 clades (Figure 3). Consistent with Harmon et al. (2010) we found very little support for EB models (only 6 clades supported EB with >75% AIC_*w*_), suggesting that “early bursts” of trait evolution are indeed be rare in comparative data (but see Slater and Pennell, 2014). Larger clades commonly had very high support for a single model (of the 101 clades consisting of more than 100 taxa, 44 had >90% of the AIC weight on a single model), and that was overwhelmingly likely to be an OU model (42/44 clades).

**Figure 3.**
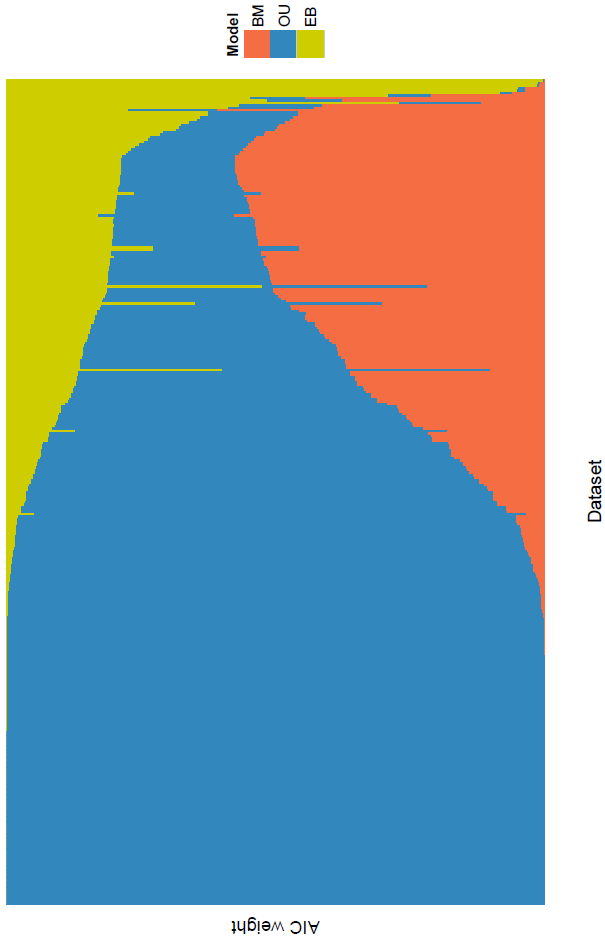
The relative support, as measured by AIC weight, for the three models used in our study (BM, OU, and EB) across all 337 datasets. An OU model is highly supported for a majority of the datasets.

We limit our analyses of model adequacy to only the most highly supported model in the candidate set, as supported by AIC. We did this to present a best-case scenario; if a model had very little relative support, it would be unremarkable if it also had poor adequacy (but see Ripplinger and Sullivan, 2010). Even considering only the best of the set, in general, the datasets often deviated from the expectations of the model in at least some ways (Figure 4). Of the 72 comparative datasets of SLA, we detected deviations from the expectations in 32 datasets (using a cutoff of *p* = 0.05), 33 by at least two, and 17 by three or more. Results were similar in the seed mass data (of the 226 seed mass datasets, we detected deviations in 153 datasets with at least one test statistic, 128 by at least two and 74 by three or more) and leaf nitrogen content (of the 39 datasets, we detected deviations in 19 by at least one, 24 by at least two, and 11 by three or more test statistic).

**Figure 4.**
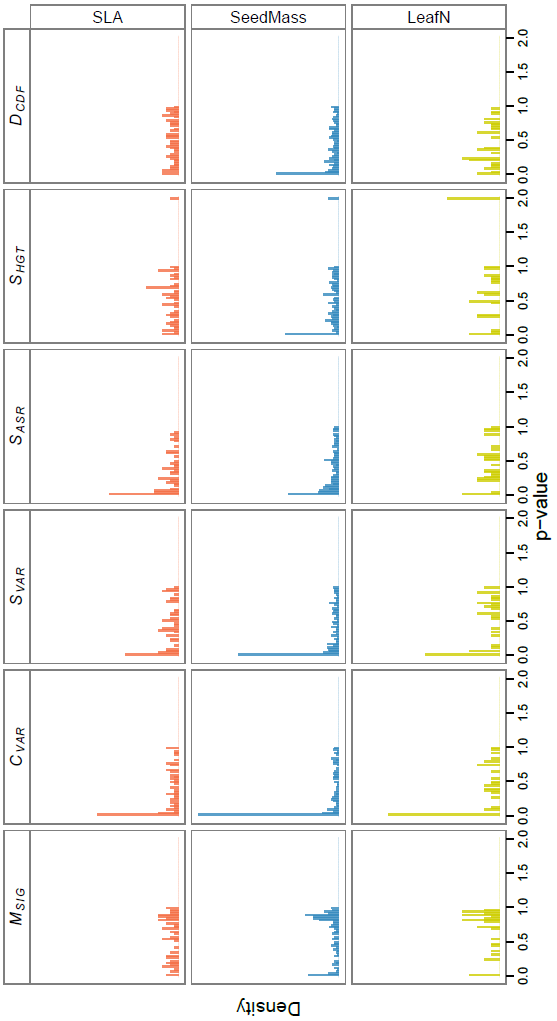
The distribution of *p*–values for our six test statistics over all 337 datasets in our study after fitting the models using ML. The *p*–values are from applying our model adequacy approach to the best supported of the three models (as evaluated with AIC). Many of the datasets deviate from the expectations under the best model along a variety of axes of variation. Deviations are particularly common for the coefficient of variation *C*_VAR_ and the slope of the contrasts against their expected variances *S*_VAR_.

Some test statistics were much more likely to detect model violations than others. In 163 cases *C*_VAR_ revealed the data deviated significantly from the expections of the best model. In 118 cases, *S*_VAR_ did. The rate of deviation was much somewhat lower for the other test statistics (*M*_SIG_: 39, *S*_ASR_: 84, *S*_HGT_: 54, *D*_CDF_: 67).

Across all 337 datasets, 133 are adequately modeled by either BM, OU or EB. As stated above, the numbers of models that showed deviations with at least one test statistic may be somewhat overinflated. However, the proportion of clades in which p-values were less than 0.05 is much, much greater than the error rates we found in our simulations. And the proportions for each individual test statistic is much higher than would be expected by chance.

As the subclades are not independent (overlapping sets of taxa are present in family, order and time-slice phylogenies), conventional statistics, such as linear regression, are not straightforward to apply across datasets. Nonetheless, the trend is clear: the larger the phylogeny, the more likely OU is to be highly supported and the more likely the model is to be inadequate. There is a strong relationship between the size of a subclade and the overall distance between observed and simulated test statistics, as measured by the Mahanalobis distance (figurFigure 5). This is not simply an artifact of conducting the analyses using a larger number of contrasts for the test statistics — if the model was adequate at all scales, there would be no relationship between the Mahalanobis distance and the size of the phylogeny. Because larger clades also tended to support a single model, the datasets for which the best model had a very poor absolute fit also had the most substantial difference between the relative fits of the three models (figure S1). There was a much weaker relationship between clade age and model adequacy (figure S2).

**Figure 5.**
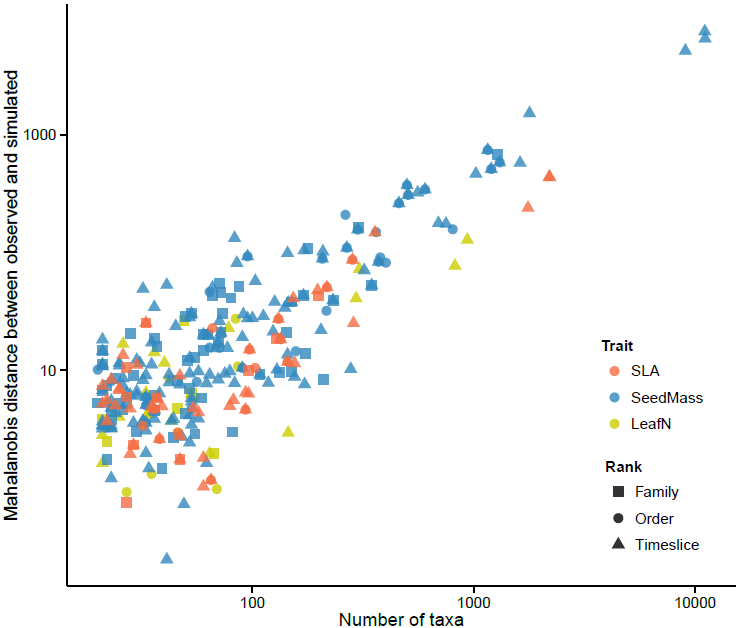
The relationship between clade size and a multivariate measure of model adequacy. The Mahalanobis distance is a scale-invariant metric that measures the distance between the observed and simulated test statistics, taking into account the covariance between test statistics. The greater the Mahalanobis distance, the worse the model captures variation in the data. Considering only the best supported model for each clade (as chosen by AIC), there is a striking relationship between the two — the larger the dataset, the worse the models performed (note the logarithmic scale). If the models were equally likely to be adequate at all scales, we would expect no relationship.

## Discussion

### Why does model adequacy matter?

Whatever inferences we want to make from comparative data — e.g., characterizing broad-scale patterns of evolution through time, investigating correlations between characters or testing hypotheses about the processes that have driven trait evolution over macroevolutionary time — it is important that our chosen statistical model captures variation in the data *relevant to the question being addressed*. If, for example, the goal is to assess variation in macroevolutionary rates over time, it is essential that the model does a good job of explaning temporally heterogeneity. If we want to know about the slope of an evolutionary allometric relationship, we need a model that provides a meaningful estimate of this parameter (Hansen and Bartoszek, 2012). Comparing the fit of a model to a set of alternatives (using likelihood ratio tests, Information Theoretic metrics, Bayes Factors, etc.) can only allow for a relative assessment of the suitability of the model for the task. Such a model comparison approach does not provide any information about whether a model will allow us to actually get at the question we are interested in.

The flipside of this is that tests of model adequacy, such as ours, are designed to measure the absolute fit but not the absolute appropriateness of the model. We know that all of the models used in comparative biology are wrong. Whether they are useful or not will depend on the question being addressed. We are far from the first to suggest that model adequacy is important to consider when using comparative methods (see, for example Felsenstein, 1985,Felsenstein 1988; Harvey and Pagel, 1991; Garland et al., 1992; Díaz-Uriarte and Garland, 1996; Hansen and Martins, 1996; Price, 1997; Garland et al., 1999; Garland and Ives, 2000; Hansen and Orzack, 2005; Hansen and Bartoszek, 2012; Felsenstein, 2012; Boettiger et al., 2012; Slater and Pennell, 2014; Beaulieu et al., 2013; Blackmon and Demuth, 2014). The contribution of our paper is to generalize many of these previous approaches into a single, flexible statistical framework.

Again, we emphasize that simply because a dataset deviates from the expectations of the model does not imply that the model should necessarily be rejected. In our analyses of model adequacy across the 337 Angiosperm clades, we were focused on whether the model was suitable for measuring rates of evolution, which is dependent on the model being a good one (Hunt, 2012). For other questions, the fact that a model fails to capture some aspects of the variation in the data may not be that important. For example, if our question was that of Harmon et al. (2010) — are early bursts of evolution common in macroevolution? — we could conclude with good certainty that they are not. Our datasets may not be well described by an OU model, but they are certainly nothing like what we would expect under an early burst scenario. Likewise, if we are interested primarily in whether there is a pattern of correlation between two traits, the fact that the model we used is not adequately describing much of the variation will in many cases, not greatly impact the qualitative conclusions.

However, we view the most interesting cases to be where the best model does not adequately describe the variation of interest. The way in which a model fails can provide a richer understanding of our data and the processes that have driven the patterns we observe (Gelman and Shalizi, 2013). First, model inadequacy can point to problems in the data. We suspect that this is likely a common cause of poor model fit. For the empirical analyses, we used a very large phylogeny of An-giosperms that was constructed to test specific global-scale biodiversity questions (Zanne et al., 2014). We recognize that the tree is poorly resolved in many places (particularly, near the tips) and is likely ill-suited for addressing more detailed, clade-specific questions (see the recent critique by Donoghue and Edwards, 2014). Specifically, the inaccurate placement of species will, on average, cause evolutionary rates to be inflated, which is precisely what we find (see below). However, we emphasize that phylogenetic error is likely ubiquitous and this problem is certainly not limited to the tree we used. Likewise, the dataset we assembled is rather heterogeneous in terms of quality; the data was originally collected for a diverse set of reasons and some groups have been measured much more carefully than others. And while we have done our best to clean the data, errors undoubtedly remain.

Second, and most excitingly, the failure of a model to adequately describe relevant aspects of the data can provide insight into the processes we have failed to consider in our model (Gelman and Shalizi, 2013). For example, if a model fails to capture variation relative to time (evaluated by the test statistic *S*_HGT_), this suggests that temporal heterogeneity has been greater than we allowed for. The causes of such heterogeneity have long been a topic of interest in macroevolutionary studies (e.g., Simpson, 1944; Foote, 1997) and there has been a great deal of recent development towards more complex rate-varying models (e.g., O’Meara et al., 2006; Thomas et al., 2006; Eastman et al., 2011; Weir and Mursleen, 2013; Rabosky et al., 2014). Likewise, failure to adequately describe variation across the clade may indicate that the existence of multiple macroevolutionary optima (sensu Hansen, 2012) are driving the dynamics of traits over time (see Hansen, 1997; Butler and King, 2004; Beaulieu et al., 2012; Ingram and Mahler, 2013; Uyeda and Harmon, 2014, for models that have been used to capture these dynamics).

Model inadequacy may also suggest types of models that have not previously been considered. For example, if recently diverged species tend to more dissimilar than can be accounted for under a simple diffusion model such as BM or OU, this may be the result of character displacement. However, almost no phylogenetic models have been put forth that explicitly model interactions between lineages (but see Nusimer and Harmon, 2014). Or if traits have lower variance than expected under an OU process, this may be the result of hard bounds. Boucher et al. (2014) recently argued that this is the case for climatic niches and that alternative models need to be developed for this case. Of course, a researcher may discover that her dataset is poorly described by all of the currently available models. Aside from deriving new models specific to her question and dataset, she should at least carefully examine the extent to which model misspecification is likely to affect the major conclusions and proceed forward with due caution.

### Implications for empirical studies

In our analysis of Angiosperm functional traits, we found common macroevolutionary models to often be poor descriptors for the patterns of variation and likely inadequate for estimating evolutionary rates. While there are certainly a number of important caveats to our analysis (discussed above), the overall trends are clear. This should certainly give researchers some pause about the models routinely used in our field — especially as they are often used in a model comparison framework to evaluate the “tempo and mode” of macroevolution. We argue that our results strongly suggest that we may often be missing a large part of the story.

The 337 comparative datasets we analyzed varied in terms of traits, size, age and placement in the Angiosperm phylogeny. Nonetheless, several general patterns emerge. An OU model, was by and large, the most supported of the three we examined. In an analysis of 67 comparative datasets consisting of size and shape data from a variety of animal taxa, Harmon et al. (Harmon et al., 2010) also found substantial support for OU models, though for their datasets, BM was more commonly chosen by AIC. (We note, however, that many of their datasets were quite small; see Slater and Pennell, 2014). Since their paper, a substantial number of studies conducted in a diverse array of groups have also found OU models to be preferred over BM models (e.g., Burbrink et al., 2012; Quintero and Wiens, 2013; LÓpez-Fernàndez et al., 2013; Thomas et al., 2014).

The tendency of OU to explain data better than BM has inspired diverse process based explanations, including stabilizing selection, evolutionary constraints and the presence of “adaptive zones” (Hansen and Martins, 1996; Butler and King, 2004; Hansen, 2012; Pennell and Harmon, 2013). If the widespread support for OU models was indeed caused by the biological processes that have been proposed, we would expect that an OU model would also be widely adequate. However, this is not what we found. The datasets deviated significantly from the distributions expected under OU models, most often detected with *C*_VAR_ and *S*_VAR_ but frequently with others as well. OU models often failed to capture other important types of heterogeneity — variation with respect to rate variation (*M*_SIG_), trait values (*S*_ASR_) and time (*S*_HGT_). Additionally, a substantial number of datasets were not well-modeled by a multivariate normal distribution (*D*_CDF_). These results suggest a statistical explanation for the high support for OU models. OU predicts higher variance near the tips of the phylogeny than do BM or EB models (see figure 1 in Harmon et al., 2010). Heterogeneous evolutionary processes, phylogenetic misestimation and measurement error could also produce such a pattern. In light of our results from model adequacy, it seems likely that OU is often supported because it is able to accommodate more “slop” (phylogenetic and trait error in addition to model misspecification) than the other models. This is not to say that the processes captured by OU models are unimportant in macroevolution, but rather that OU models may be favored for reasons that are more statistical than biological. Future, and hopefully more widely adequate, models of trait evolution could be developed that both include aspects of the OU model, especially the bounds on trait values, while incorporating additional biological realism (for a recent example of such a model, see Nusimer and Harmon, 2014).

The way in which the observed test statistics deviate from the simulated values also supports the claim that the widespread support for OU is largely a statistical artifact. Model violations were most frequently detected by the variance estimate, CVAR. If the evolutionary process (or, alternatively, phylogenetic/measurement error) is heterogeneous across the tree, the lineages in some parts of the clade will be much more divergent than in others. The only way for the model to account for the highly divergent groups is to estimate a large *σ*^2^ (and/or a small *α* parameter for the OU model). The unit tree formed by these parameter estimates will have long branches across the entire tree. In the less divergent parts of the tree, the contrasts calculated on this unit tree will be small, relative to what we expect under BM. So perhaps counter-intuitively, when heterogeneity in processes across taxa cause the estimated global rates of divergence to be inflated, resulting in a higher value for *C*_VAR_.

The second major take-home from the empirical analyses is that error, both in trait values and phylogenies, can have serious consequences for model adequacy. We frequently detected deviations from model expectations with *S*_VAR_, the slope between the contrasts and their expected variances. This is indicative the rate of evolution appears to be varying with regards to branch length over which it is measured. This seems unlikely to be attributable to any biological process; it is far more probable that this reflects phylogenetic error (particularly, branch length error). Above, we outlined some of the deficiencies of the datasets we used in this paper but argue that these are likely to be widespread in comparative data. The test statistics outlined above can serve as useful diagnostics to aid researchers in identifying outliers that may be driving the pattern. We recommend that researchers faced with an inadequate model plot the magnitude of the contrasts on to the unit tree; this will usually be much more informative with regards to the model fit than plotting the magnitude of the contrasts on the original phylogeny. Exceptionally large or small contrasts on the unit tree can provide clues as to where the data may be erroneous. If phylogenetic error were causing poor model fits, we would predict that many of the anomalous contrasts would occur in parts of the tree that are poorly supported.

### Extensions of our approach

There are a number of additional ways our approach could be extended. First, we have only considered a limited set of test statistics. We chose them because each of these has a clear statistical expectation and observed deviations from them have intuitive biological explanations. However, they are certainly a subset of all possible test statistics that could be applied. For example, because contrasts are i.i.d., there should be no autocorrelation between neighboring contrasts; the test statistics could be expanded to detect non-zero autocorrelation. Second, as stated above, our approach can be applied equally well to phylogenetic regression models, such as phylogenetic generalized least squares (Grafen, 1989; Martins and Hansen, 1997) or phylogenetic mixed models (Lynch, 1991; Housworth et al., 2004; Hadfield and Nakagawa, 2010), where concerns regarding model adequacy are just as pertinent (Hansen and Bartoszek, 2012). While our approach can be used to assess the adequacy of the phylogenetic component of regression models “out of the box”, additional steps are required to assess the adequacy of the linear component. Third, our method was designed for quantitative trait models that assume data can be modeled with a multivariate normal distribution. We need general model adequacy approaches for other types of traits, such as: discrete traits (i.e., binary, multistate, ordinal; see Beaulieu et al., 2013; Blackmon and Demuth, 2014; Maddison and FitzJohn, 2014, for recent discussions of this); traits that influence speciation rates (e.g., Maddison et al., 2007; FitzJohn, 2010) and quantitative trait models that do not predict a multivariate normal distribution of traits (Landis et al., 2013; Schraiber and Landis, 2014).

It may also be possible to extend our approach with an eye towards model selection. Slater and Pennell (2014) developed their posterior predictive simulation approach (which is related to our method) to distinguish between a BM model and one where rates of evolution decreased through time. They chose test statistics specifically to address this question. Slater and Pennell found using posterior predictive fit as a model selection criterion to be much more powerful than comparing models using AIC or likelihood ratio tests, particularly when “outlier taxa” (lineages where the pattern of evolution deviates from the overall model) were included in the analysis. The logic of Slater and Pennell could be extended to other scenarios; to test some evolutionary hypotheses, we may care a lot about whether a model explains varation along some axes but be less concerned about others. This is a question-specific approach to model selection and has been developed in the context of molecular phylogenetics (Bollback, 2002; Lewis et al., 2014). This is also the essence of the Decision-Theoretic approach to model selection (Robert, 2007), which has also been well-used in phylogenetics (Minin et al., 2003), but has not previously been considered in PCMs.

## Arbutus

We have implemented our approach in a new R package arbutus. It is available on github https://github.com/mwpennell/arbutus. For this project, we have also adopted code from the ape (Paradis et al., 2004), `geiger` (Pennell et al., 2014) and `diversitree` (FitzJohn, 2012) libraries. We have written functions to parse the output of a number of different programs for fitting trait evolution models (see the `arbutus` website for an up-to-date list of supported models and packages). As this approach was developed to be general, we have written the code in such a way that users can include their own test statistics and trait models in the analyses.

## Concluding remarks

Attempts to assess the adequacy of phylogenetic models are almost as old as modern comparative phylogenetic biology. In the 1980s and 1990s much discussion surrounded the appropriateness of various methods and models (Felsenstein, 1985,Felsenstein 1988; Harvey and Pagel, 1991; Garland et al., 1992; Díaz-Uriarte and Garland, 1996; Price, 1997; Garland et al., 1999; Garland and Ives, 2000). We argue that this discussion is key to progressing in our field. This is not simply because we are concerned that many inferences may not be robust to model violations. Rather, we believe that considering model adequacy can help suggest new ways of thinking about how to combine data and models to test macroevolutionary hypotheses.

## Acknowledgments

We would like to thank the members of the Tempo and Mode of Trait Evolution Working Group at the National Evolutionary Synthesis Center (NESCent) for their suggestions and encouragement. We thank Josef Uyeda, Daniel Caetano, Will Pearse and Sally Otto for their thoughtful comments on the manuscript. Paul Joyce, Graham Slater, Jeremy Brown, and Cécile Ané also provided valuable insights into this project. This manuscript greatly benefitted from the comments of Troy Day, Scott Steppan, and two anonymous reviewers. Last, we are grateful to the researchers who made their data available; this project would not have been possible without it. MWP was supported by a NESCent graduate fellowship and a NSERC postgraduate fellowship. This work was also supported by NSF grants awarded to LJH (DEB 0919499 and 1208912).

## Results from Bayesian analyses

As with the likelihood results (described in main text), OU models were highly supported across many datasets; 177/337 clades had the highest DIC weight (DIC*_w_*) on an OU model; 156 of them with greater than 75% of the total DIC*_w_* (see Figure S3). While a generally similar pattern of model support holds for both likelihood and Bayesian inference, the likelihood analyses are much cleaner (compare figure 3 and figure S3). This differnce can be explained by the fact that there is a tight statistical relationship between the AIC values for these three models. If two models have identical likelihoods, the AIC scores, defined as 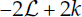 (where 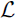 is the log-likelihood of the model and *k* is the number of parameters) will differ by 2. As BM is a special case of both OU and EB, in opposite directions in model space, the highest AIC*_w_* possible for BM is ~0.731. The rare clades where both OU and EB have higher support than BM likely reflect problems in optimization. Calculating DIC values from posterior samples is inherently more stochastic; if there is little information in data, the best DIC model will depend on the values sampled by the chain.

For the model adequacy results, the results were also very similar to that of the likelihood analyses (compare to Results section in the main text). The adequacy of these simple models was poor across the majority of the datasets (figure S4). Again, we limit our analyses of model adequacy to only the most highly supported model in the candidate set.

Of the 72 comparative datasets of SLA, we detected deviations from the expectations of the best supported model using at least one test statistic in 35 cases, 26 by at least two, and 19 by three or more. For the seed mass data, we detected deviations with at least one test statistic in 173 cases (by two or more in 109 datasets and by at least three in 72 cases). 24/39 leaf nitrogen datasets were found to be inadequately described by the best supported model with at least one test statistic (13 by at least two and 10 by at least three).

Also, similar to the likelihood analyses, the frequency at which deviations were found differed between the test statistics. In 171 cases, we detected model misspecification with *C*_VAR_ and with *S*_VAR_, 141 (*M*_SIG_: 24, *S*_ASR_: 101, *S*_HGT_: 78, *D*_CDF_: 67). Again, only 105 datasets were adequately modeled by one of the three models in our candidate set. There was a strong relationship between model (in)adequacy and clade size (figure S5), but less so for clade age (figure S6).

**Figure S1.**
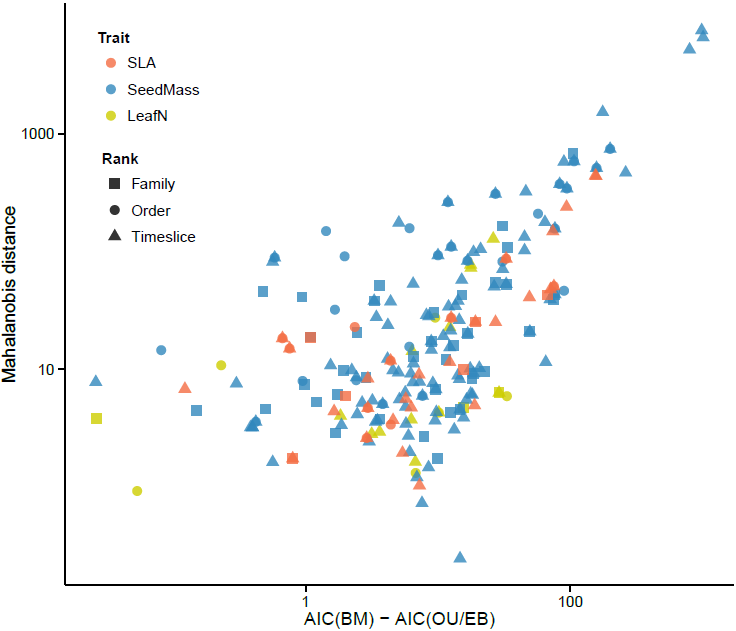
The relationship between relative and absolute fit. For every clade for which a more complex model (OU, EB) was favored over BM using AIC, the Mahalanobis distance between the observed test and simulated test statistics is plotted against the improvement in AIC for the more complex model compared to BM. (Note that as all AIC values were negative, larger differences mean greater relative support). The greater the relative fit of a more complex model, the more likely the model was to be inadequate. This result in primarily driven by clade size but serves to emphasize the distinction between relative and absolute fit.

**Figure S2.**
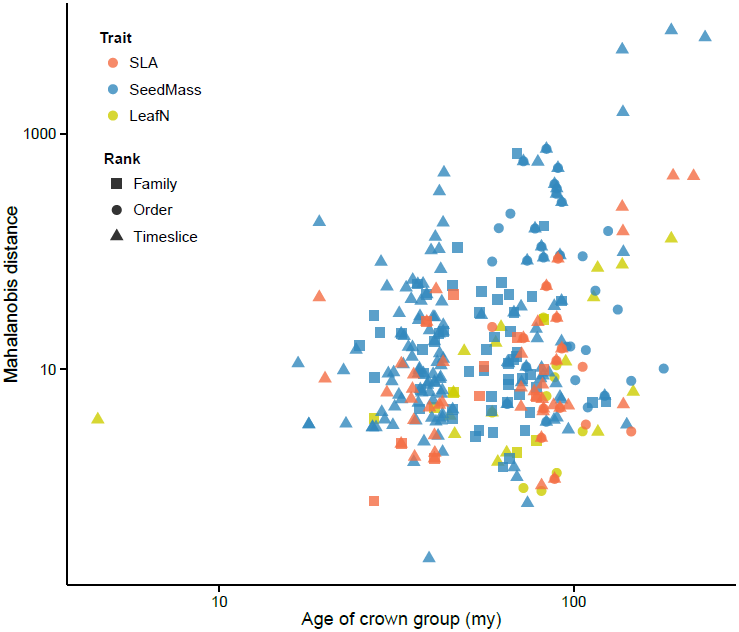
The relationship between clade age and a multivariate measure of model adequacy. Considering only the best supported of the three models (as selected by AIC, after fitting the models using ML), there is no apparent relationship between the age of clade and the distance of the observed and simulated test statistics, as measured by the Mahalanobis distance. Contrast this figure with figure 5, which demonstrates a very tight relationship between clade size and model inadequacy.

**Figure S3.**
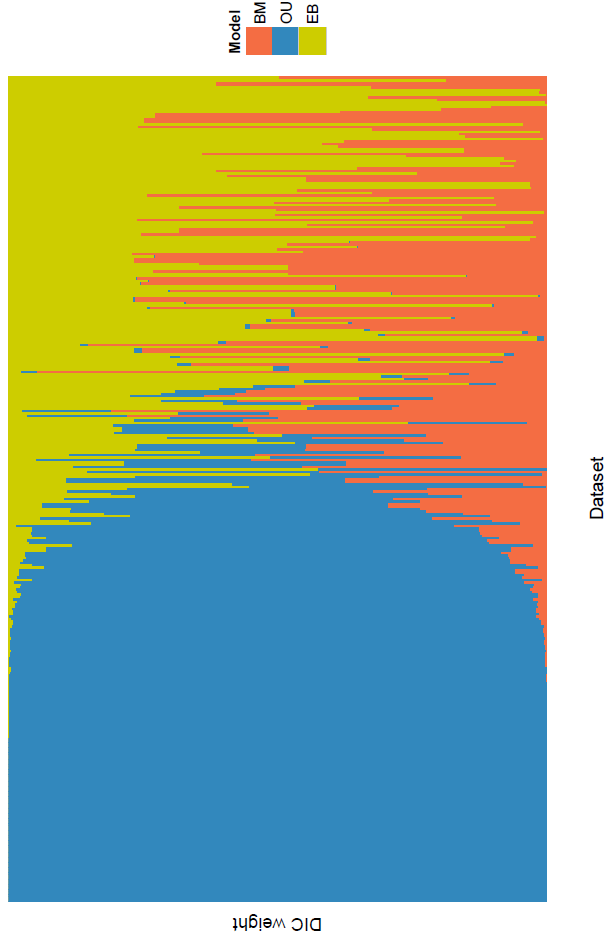
The relative support, as measured by DIC weight, for the three models used in our study (BM, OU, and EB) across all 337 datasets. All models were fit with MCMC. Like the model comparisons done with AIC, an OU model is highly supported for a majority of the datasets.

**Figure S4.**
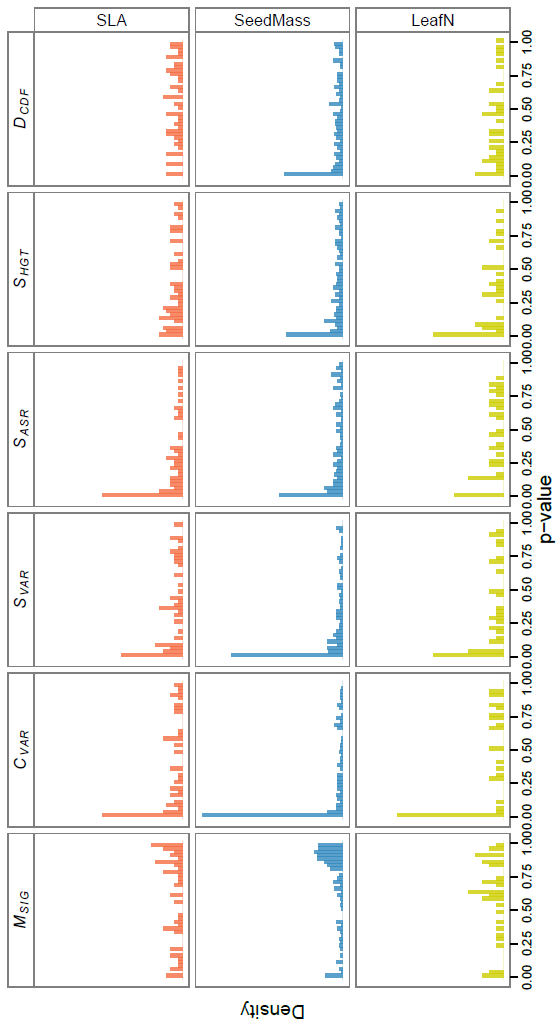
The distribution of *p*–values for our six test statistics over all 337 datasets in our study after fitting the models using MCMC. The *p*–values are from applying our model adequacy approach to the best supported of the three models (as evaluated with DIC). Many of the datasets deviate from the expectations under the best model along a variety of axes of variation. Deviations are particularly common for the coefficient of variation *C*_VAR_ and the slope of the contrasts against their expected variances *S*_VAR_.

**Figure S5.**
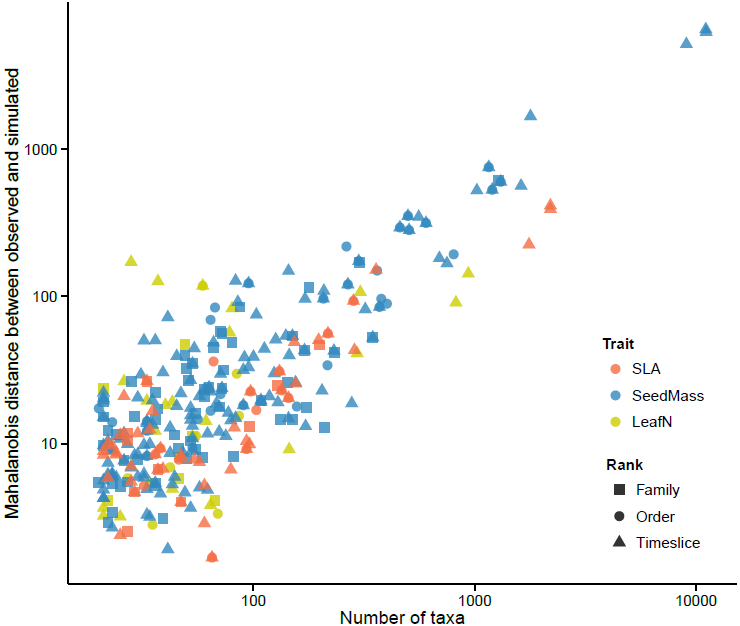
The relationship between clade size and a multivariate measure of model adequacy from the Bayesian analysis. The Mahalanobis distance is a scale-invariant metric that measures the distance between the observed and simulated test statistics, taking into account the covariance between test statistics. The greater the Mahalanobis distance, the worse the model captures variation in the data. Considering only the best supported model for each clade (as chosen by DIC), there is a striking relationship between the two — the larger the dataset, the worse the models performed (note the logarithmic scale). If the models were equally likely to be adequate at all scales, we would expect no relationship.

**Figure S6.**
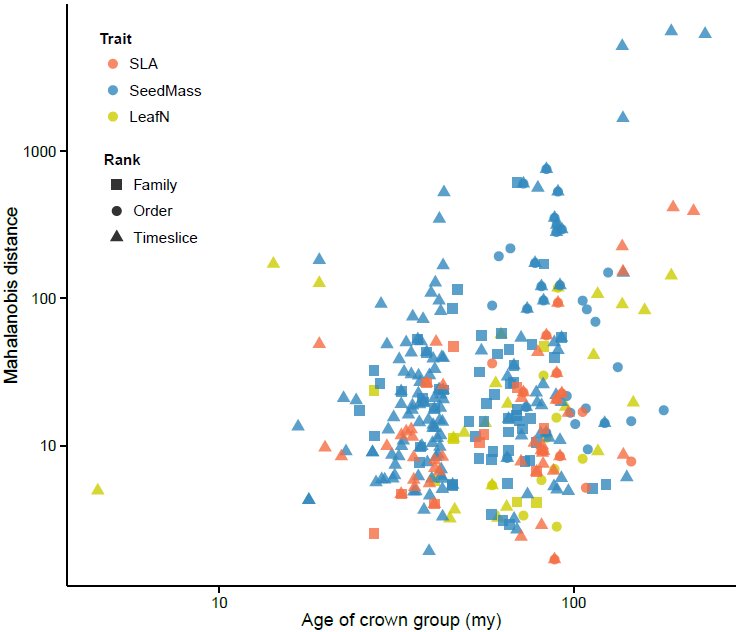
The relationship between clade age and a multivariate measure of model adequacy. Considering only the best supported of the three models (as selected by AIC, after fitting the models using MCMC), there is no apparent relationship between the age of clade and the distance of the observed and simulated test statistics, as measured by the Mahalanobis distance. Contrast this figure with figure S5, which demonstrates a very tight relationship between clade size and model inadequacy.

**Figure S7.**
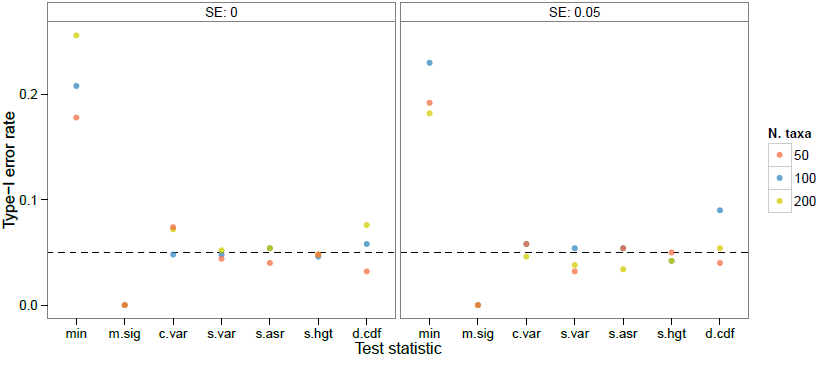
Type-1 error rates for data simulated under a Brownian motion (BM) model. We simulated 500 datasets under for 3 different tree sizes (N = {50,100,200}, represented by the different colors) and two known values of standard error (0 and 0.05, left and right panel, respectively). The Type-1 error rates for each test statistic are consistently around or lower than 0.05 threshold. However, the frequency at which *at least one* of the test statistics deviated significantly from the expectations (the variable “min” on the left side of each plot) was substantially greater, rising to above 0.2% in some cases. See text for why we decided against correcting for the effect of multiple comparisons in the analysis.

**Figure S8.**
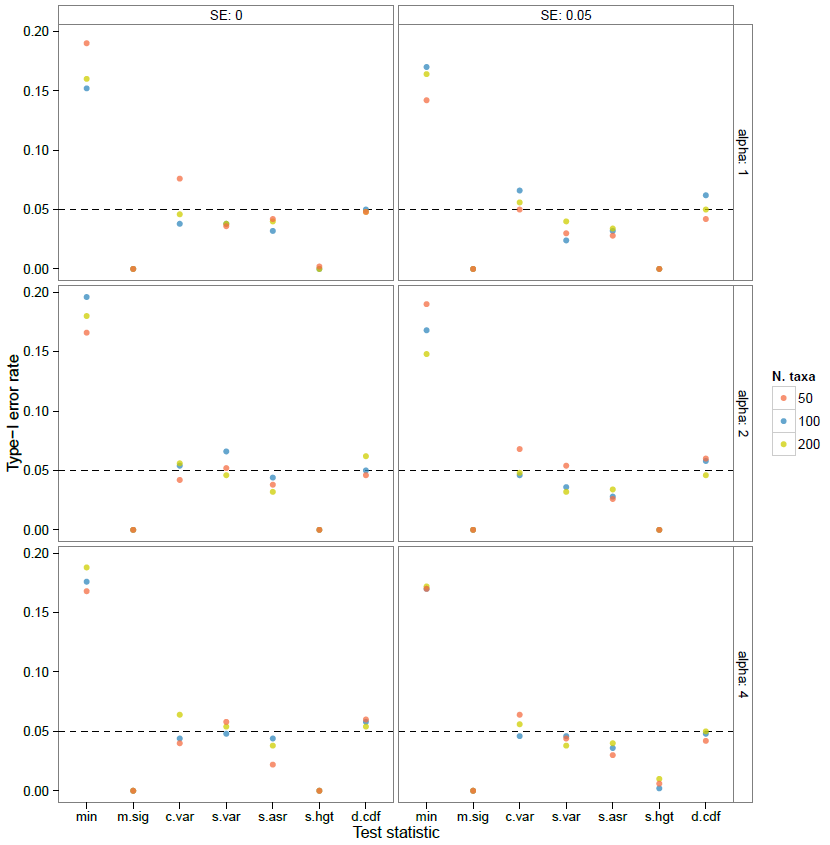
Type-1 error rates for data simulated under an Ornstein-Uhlenbeck (OU) model. We simulated 500 datasets under for 3 different tree sizes (N = {50,100,200}, represented by the different colors) and two known values of standard error (0 and 0.05, left and right panel, respectively). We also simulated under three values for the *α* parameter (*α* = {1,2,4}, top, middle and bottom panel), representing phylogenetic half-lives of 69%, 35%, 17% of total tree depth respectively The Type-1 error rates for each test statistic are consistently around or lower than 0.05 threshold. However, the frequency at which *at least one* of the test statistics deviated significantly from the expectations (the variable “min” on the left side of each plot) was substantially greater, approaching 0.2% in some cases. See text for why we decided against correcting for the effect of multiple comparisons in the analysis.

**Figure S9.**
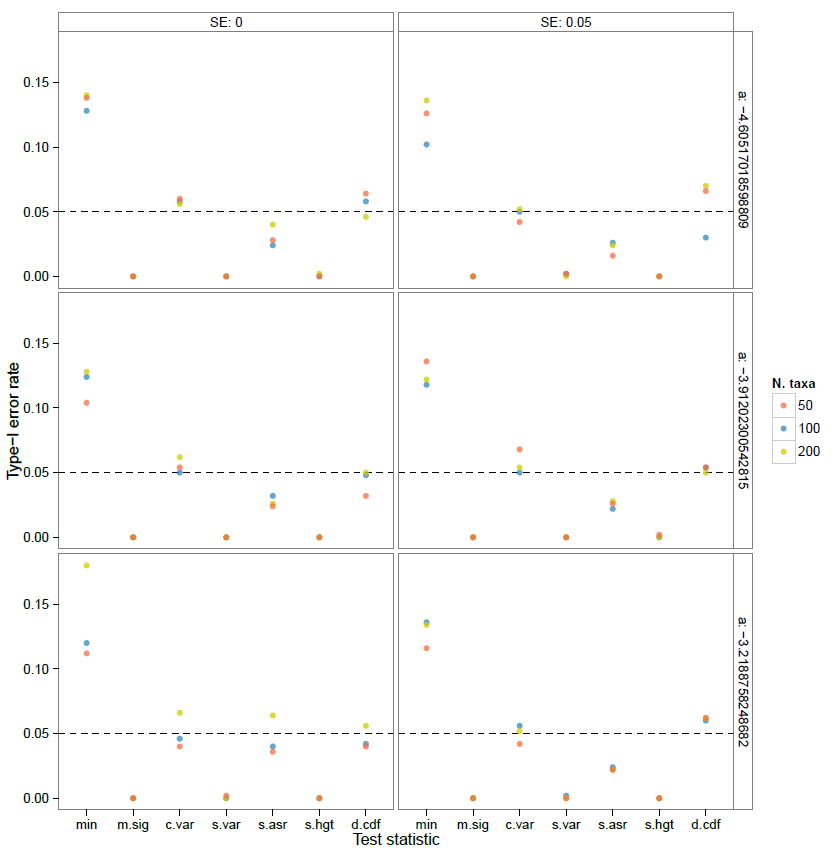
Type-1 error rates for data simulated under an Ornstein-Uhlenbeck (OU) model. We simulated 500 datasets under for 3 different tree sizes (N = {50,100,200}, represented by the different colors) and two known values of standard error (0 and 0.05, left and right panel, respectively). We also simulated under three values for the exponential rate of slowdown, *a* (*a* = {log(0.01),log(0.02),log(0.04)}, top, middle and bottom panel), which translate to the rate of trait evolution halfing every 0.15, 0.17, and 0.21 time units, respectively (note that the tree was scaled so the total depth was equal to unity). The Type-1 error rates for each test statistic are consistently around or lower than 0.05 threshold. However, the frequency at which at least one of the test statistics deviated significantly from the expectations (the variable “min” on the left side of each plot) was substantially greater, approaching 0.15% in some cases. See text for why we decided against correcting for the effect of multiple comparisons in the analysis.

